# The global groundwater resistome: core antibiotic resistance genes, their dynamics and drivers

**DOI:** 10.1101/2022.11.14.516424

**Authors:** Ioannis D. Kampouris, Johan Bengtsson-Palme, Doreen Babin, Sara Gallego, Bing Li, Kornelia Smalla, Thomas U. Berendonk, Uli Klümper

## Abstract

Despite the importance of groundwater as a drinking water resource, currently, no comprehensive picture regarding the global levels of antibiotic resistance genes (ARGs) in groundwater environments exists. Moreover, the biotic and abiotic factors that shape the groundwater resistome on the global scale remain to be explored. Herein, we attempted to fill this knowledge gap through *in silico* re-analysis of publicly available global groundwater metagenomes. First, nine ARGs encoding resistance to aminoglycosides (*aad*A, *aph*(3’), and *ant*(3’’)), sulfonamides (*sul*1 and *sul*2), β-lactams (*bla*_OXA_ and *bla*_TEM_), tetracyclines (*tet*(C)) and macrolides (*msr*(E)) were identified to constitute the core groundwater resistome with high detection and abundance levels. Second, the global drivers of groundwater resistome composition were identified by applying a structural equation model with mixed effects to disentangle the individual contributions of each abiotic and biotic factor. Most notably, global effects of the origin of groundwater samples on the resistome were detected with samples from high-income countries (HICs) constantly displaying lower ARG and mobile genetic element (MGE) abundances than those from low-and-middle-income countries (LMICs). While these effects were consistent across antibiotic classes, biotic factors such as interactions of the groundwater microbiome with fungal or bacterial natural producers of antibiotics, or the co-occurrence of ARGs on mobile genetic elements (MGEs) played significant roles in shaping abundance patterns of resistance towards individual antibiotic classes. Only few ARGs correlated to individual bacterial genera, with microbial community composition in general weakly associated with resistome composition. In conclusion, we provide a first global picture of the resistome of low-anthropogenic impacted groundwater environments and the underlying anthropogenic and biotic drivers shaping it, which can be used as a baseline in future surveillance of antibiotic resistance.

## 1. Introduction

The global rise in antimicrobial resistance (AMR) represents a major threat to human health (Laxminarayan et al., 2013). Tackling it requires a “One Health” approach that considers AMR dynamics and proliferation between the interconnected human, veterinary, and environmental spheres (Hernando-Amado et al., 2019). Within this context, drinking water resources provide one of the immediate connections between environmental and human microbiomes (Vaz-Moreira et al., 2014). Among these, groundwater ecosystems constitute the most common freshwater and drinking water resource in the majority of the world (Griebler and Avramov, 2015; Herrmann et al., 2019; Szekeres et al., 2018). Groundwater environments are characterized by high microbial diversity and complexity (Flynn et al., 2013; Griebler and Avramov, 2015; Griebler and Lueders, 2009), with groundwater microbiota playing important roles in multiple biogeochemical cycles (Flynn et al., 2013; Retter et al., 2021; Sonthiphand et al., 2019). Due to their role as a major drinking water resource, understanding the spread of AMR in and through groundwater environments is highly relevant for tackling AMR through the “One Health” approach (Hernando-Amado et al., 2019). AMR in groundwaters has only been addressed in a few studies using qPCR or metagenomic approaches aiming to elucidate the occurrence of antibiotic resistance genes (ARGs) in groundwater (e.g., Szekeres et al., 2018; Zaouri et al., 2020; Zhang et al., 2019).

A global and comprehensive picture of ARG levels in groundwater, which identifies core groundwater ARGs and allows investigation of the underlying anthropogenic or non-anthropogenic factors shaping groundwater resistomes, is currently missing and could provide a baseline for future AMR surveillance efforts (Bengtsson-Palme et al., 2023). Such a global picture regarding AMR has been previously created for other environments of low anthropogenic impact, such as terrestrial soils (Bahram et al., 2018). Here, on the biotic side, a general link between ARG levels and interactions between fungal and bacterial taxa was detected (Bahram et al., 2018). Fungi can manipulate and shape the indigenous bacterial communities by producing and exuding chemical compounds (Johnston et al., 2016), including β-lactam antibiotics (e.g., penicillin) (Aly et al., 2011). Consequently, fungi-bacteria interactions have been proposed as the underlying cause of the natural prevalence of β-lactam ARGs in the environment (e.g., the occurrence of *bla*_TEM_ and *bla*_CTX-M_ variants in pristine soils) (Gatica et al., 2015). Such biotic interactions could also come into play in groundwater environments where fungi occur at relatively high abundances, mainly functioning as saprophytic organisms that enable the degradation of organic matter and organic carbon recycling (Nawaz et al., 2018). However, fungal-bacterial interactions and interactions of the groundwater microbiome with antibiotic producers could similarly affect the resistome in the relatively isolated, dark, and pristine groundwater environments.

On the abiotic side, potential anthropogenic impacts on AMR were demonstrated in groundwater reservoirs located beneath commercially operated agricultural fields irrigated with treated wastewater (Kampouris et al., 2022). Here the abundance of ARGs increased with the infiltration of antibiotics and other pharmaceuticals into the groundwater environment. However, how abiotic factors shape ARG dynamics in the majority of less-impacted global groundwater environments remains poorly understood. Apart from wastewater irrigation, general anthropogenic activities (e.g., agriculture) might affect the introduction, dissemination, and selection dynamics of ARGs in groundwater environments (Heuer et al., 2011). Regularly, high-income countries (HIC) can afford and emphasize stricter environmental and health standards, restrictions on antibiotic consumption, and sustainable agricultural practices compared to low-and-middle-income countries (LMIC) (Hosain et al., 2021). We propose that these differences could be one of the factors leading to elevated dissemination of ARGs to groundwater environments in LMICs compared to the HICs.

With the broad availability of groundwater metagenomes from various locations around the world, it has now become possible to fill these knowledge gaps and obtain a global and comprehensive picture of groundwater resistomes. In this study, we aim to explore the composition and drivers shaping groundwater resistomes through *in silico* re-analysis of publicly available global groundwater metagenomes retrieved from the NCBI sequencing read archive (SRA). We specifically identify core groundwater ARGs and explore the underlying anthropogenic and non-anthropogenic factors shaping groundwater resistomes.

## 2. Methodology

### 2.1 Data collection of groundwater metagenomes

Public metagenome datasets sampled from global groundwater environments were searched and collected from the sequencing read archive (SRA). The search queries included the terms “groundwater”, “aquifer”, or “subsurface water”, for matrices; and “shotgun sequencing” or “wgs” for the sequencing method. We mainly focused on Illumina sequencing methods, since there was no high availability of long-read sequencing datasets that would enable robust comparisons. The information from the SRA was linked to specific publications and locations, whenever available. Further, associated metadata such as geographic location, and depth were collected. The metagenome sampled from different countries were classified into two groups based on the average income, as estimated by the gross domestic product per capita, to HICs and LMICs. This division was performed on the basis that HICs have regularly stricter health and environmental regulations, with improved criteria for wastewater treatment and waste disposal.

A total of 100 metagenomic datasets retrieved from the SRA passed the initial quality criteria of sequencing reads above 90 bp and error rate <0.1. An additional approximately 30 identified metagenomes in the literature from peer-reviewed studies were unfortunately not made publicly available or did not display a high enough number of reads passing the quality criteria and were thereby also excluded from the study. For each metagenomic dataset that passed initial criteria, general quality control and trimming were performed with the tool cutadapt (v3.1, (Martin, 2011) with the command: cutadapt --cores=10 --cut 20 -q 10 --minimum-length 90 --max-n 0 -- max-ee 0.1. Setting the maximum expected error to 0.1 allowed only high-quality sequences to pass. Sequences with a length of less than 90 bp were filtered out, to ensure a sufficient read length for subsequent ARG annotation. We then used sequencing depth of reads annotated as small subunits of rRNA (SSUs) and Good’s coverage ([1-singleton_SSUs/total_SSUs]*100) for bacterial community estimation of sequencing performance and excluded samples with less than 5000 bacterial SSU reads and 95% Good’s coverage due to poor sequencing depth that could heavily compromise our analysis. This reduced our dataset to 68 samples, with 29 from HIC and 39 samples from LMIC countries. Accession numbers, metadata, and linked publications for all retrieved high-quality groundwater metagenomic datasets are given in Table S1.

### 2.2 Annotation of antibiotic resistance gene profile

To annotate ARGs in the resulting metagenomic datasets ResFinder (Version 4), a database of mobile, acquired ARGs (Bortolaia et al., 2020), was translated from nucleotide sequences into amino acid sequences using Biopython (Cock et al., 2009). ARGs were annotated against the translated ResFinder database using the command “blastx” in DIAMOND (Buchfink et al., 2014) with the following parameters: minimum identity 99%, minimum match length 30 amino acids. The parameters were chosen to be conservative to reduce false positive hits. In the case of paired-end sequencing, matches on the second paired read were counted only if there was no match on the first read. Since assembly into contigs might result in the loss of low-abundant ARGs that are not part of assembled contigs (Abramova et al., 2023), short-read-based analysis against a curated database was selected for inferring abundances, as this enables higher sensitivity (Abramova et al., 2023). Assembly was performed only in the context of finding potential co-localizations.

### 2.3 Annotation of selected mobile genetic elements

To annotate selected mobile genetic elements (MGEs) we used the already prepared Mobile-OG database of proteins that associate with MGEs (Brown et al., 2022) combined with a more conservative database (Pärnänen et al., 2018). From all annotated MGE markers, we selected Class 1 integron integrase (*intI*1)*, kor*B (as a proxy for IncP-1 plasmids), *TnpA* transposase (*tnp*A), and total phage abundance, all associated with the capturing and integration of ARGs (Brown-Jaque et al., 2015; Gillings et al., 2015; Linares et al., 2006; Schlüter et al., 2007; Sommermann et al., 2018) for subsequent analysis.

### 2.4 Assembly of metagenomes for identifying co-localized genes

To identify which ARGs are located on MGEs, we performed the assembly of groundwater metagenomes into contigs using MEGAHIT (Li et al., 2015). ARGs and MGEs were annotated via DIAMOND, using the same methodology for read-based analysis. The minimum aligned region for assembly-based analysis had a minimum alignment length of 100 amino acids (300 bp). The results of co-localization were visualized via “ggvenn” (Linlin Yan, 2023) and “ggsankey”(David Sjoberg, 2023).

### 2.5 Annotation of taxonomical composition

The tool METAXA2 (version 2.2.3) (Bengtsson-Palme et al., 2015) was used for the identification of total small subunits of the ribosome (SSU) 16S rRNA and 18S rRNA genes for prokaryotes and eukaryotes, respectively, from metagenomic datasets to determine phylogenetic composition, using the default settings. In addition, we screened for crAssphage sequences, a viral indicator for anthropogenic fecal pollution (Karkman et al., 2019), with “ngless” (Coelho et al., 2019), which utilizes a version of the BWA-MEM algorithm for alignment (Li, 2013; Li and Durbin, 2010).

### 2.6 Data analyses and statistics

Following ARG-, MGE-, and phylogenetic annotations, results were analyzed in R (R Core Team, 2021). Total bacterial and fungal counts for each metagenomic sample were calculated using the “tidyverse” packages (v1.0.4, Wickham et al., 2019). The ARG, bacterial and fungal relative abundances were calculated accordingly using “dplyr” (v.1.0.10,, Wickham et al., 2019) and “ggplot2” (Hadley Wickham, 2016).

Differences in the ARG composition based on Euclidean distance were visualized and evaluated using the “vegan” package (v.2.5.6, Oksanen J, et al., 2022) by generation of NMDS plots and statistical PERMANOVA tests. ARG and fungal or phage relative abundance were calculated as ARG reads in relation to million SSU (parts per million, ppm). All data was log_10_-transformed and pseudocounts (+1) were added to counter zeros prior to transformation. For differences in bacterial community composition, we used Bray-Curtis dissimilarity of bacterial taxa at the genus level. Unclassified genera were grouped based on their lowest classified rank.

For comparing the differential abundance of single ARGs, Wilcoxon rank-sum tests were performed. For bacterial taxa, random forest was performed to select the 20 most important taxonomic groups that contribute to the difference between HICs and LMICs with subsequent Wilcoxon rank-sum tests applied. To estimate the association of bacterial community composition with ARG dissemination, we performed Mantel tests and co-occurrence networks via Spearman rank correlation, which was visualized with the packages “igraph” (Csárdi et al., 2023) and “ggnetwork” (François Briatte et al., 2024). In addition, we performed correlation analysis of ARG abundance with the main taxonomic groups of antibiotic-producing microorganisms: a) *Actinobacteria* SSU relative abundance and b) fungal/bacterial SSU ratios using Spearman correlation and linear mixed effect models (package “lme4”, v. 1.1-3.5.2, Bates et al., 2015).

To decipher the best predictors for ARG dissemination in the groundwater environments a structural equation model was fitted to determine the combined effect of the individually identified biotic and abiotic factors on individual antibiotic classes and correlations between different antibiotic classes. Bacterial community composition was included by using the principal component that explained most of the variance (20.8%) following a PCoA analysis of the Bray-Curtis dissimilarity. This model included the different studies as a random effect, to account for technical biases between different studies. The structural equation model was calculated with the package “lavaan” (Rosseel, 2012). Only significant relationships with *p*<0.05 after correction for multiple testing are shown in the structural equation model. The results were visualized with the packages “igraph” and “ggnetwork” (Csárdi et al., 2023; François Briatte et al., 2024)

## 3. Results

### 3.1 High-quality global groundwater metagenomes for re-analysis

Out of the >100 retrieved and screened groundwater metagenomes (Table S1), 100 exceeded the initial sequencing reporting criteria (read length >90 bp, expected error rate <0.1/read) for subsequent re-analysis. From these 100 groundwater metagenomes, a further 32 metagenomes were excluded, either due to the low coverage (below 95% of bacterial SSUs) or low sequencing depth (below 5000 bacterial SSU hits) of the bacterial community metagenomes (Fig. 1A). To avoid any groundwater samples immediately polluted by fecal contamination, we screened the metagenomes for the abundance of crAssphage, a common fecal contamination indicator. Throughout all metagenomes, no crAssphage hits were detected. This resulted ultimately in a selection of 68 high-quality groundwater metagenomic samples for subsequent analysis representing diverse geographical locations with different average income levels (e.g., USA, Saudi Arabia, Japan, Germany, China, etc.).

**Figure 1.**
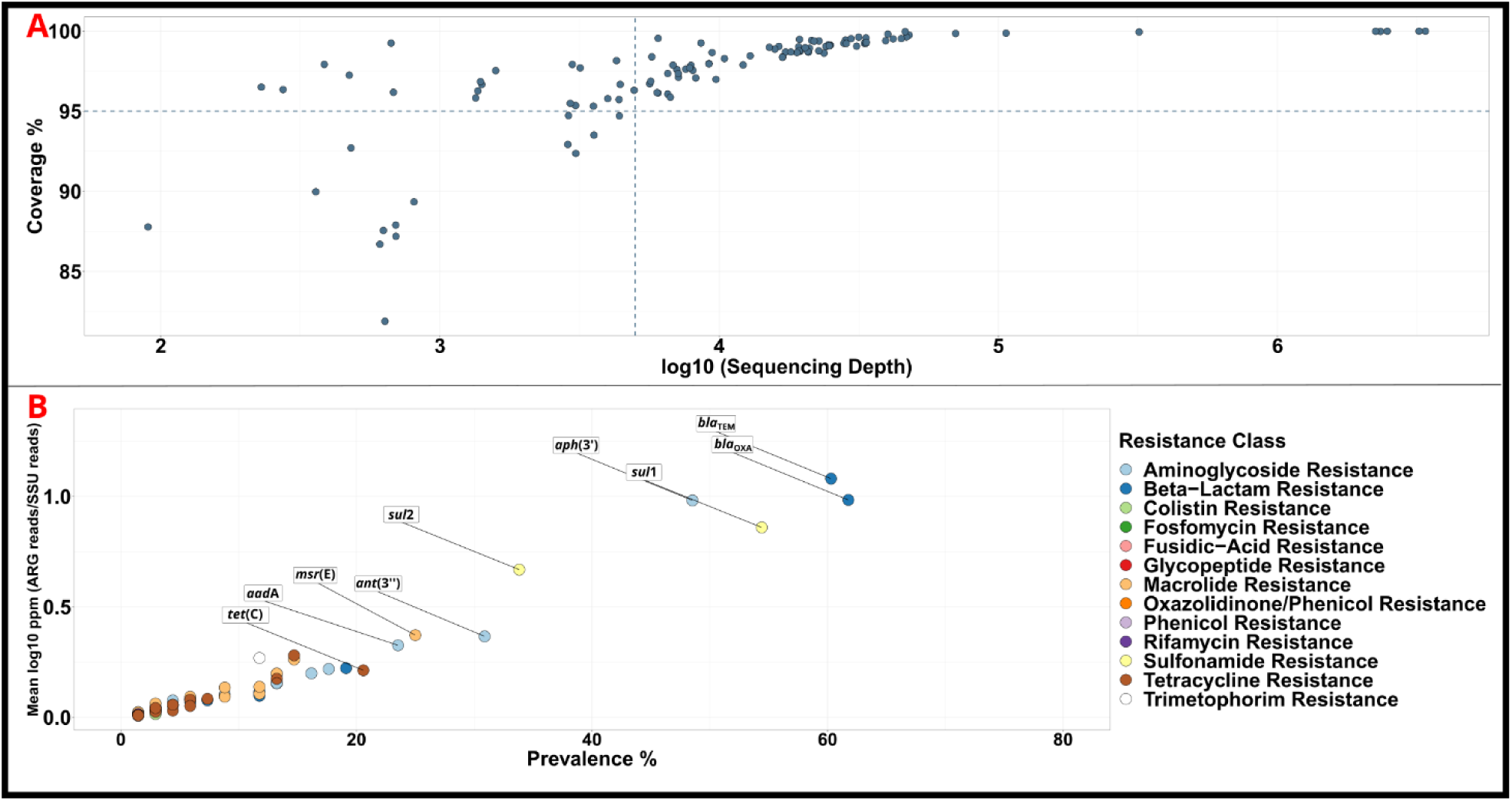
A) Good’s coverage for estimating whether the shotgun sequencing method covered at least 95% of the diversity of sampled groundwater environments at a sequencing depth of ≥5000 16S rRNA gene sequences. Only samples in the top right quadrant were included in subsequent analysis. B) The mean abundance (average in samples where ARG was detected) and the prevalence (% ARGs in detected samples) of different ARGs in the analyzed publicly available groundwater samples to identify the core groundwater resistome.

### 3.2 The core groundwater resistome: antibiotic resistance genes with the highest prevalence

Overall, 85 individual ARGs were detected across the groundwater samples as part of the global groundwater resistome, with ARGs being detected in 66 of the 68 high-quality groundwater metagenomes (Table S2). These 85 ARGs confer resistance to 13 antibiotic classes. Among these antibiotic classes, only ARGs that confer resistance to sulfonamides, macrolides, aminoglycosides, β-lactams, tetracyclines and trimethoprim were detected in at least 20% of samples and throughout, when detected, at higher relative average abundance compared to the remaining classes (Fig. 1B). On the individual ARG level, the sulfonamide ARGs *sul*1 and *sul*2, the β-lactam ARGs *bla*_OXA_ and *bla*_TEM,_ the aminoglycoside ARGs *aad*A, *aph*(3’) and *ant*(3’’), the tetracycline gene *tet*(C) and the macrolide gene *msr*(E) occurred in at least 20% of the metagenomes and were also most highly abundant on average (Fig. 1B & 2C). They hence constitute the core groundwater resistome. Using these core groundwater ARGs we subsequently investigated the underlying factors and variables that might affect their global distribution patterns.

### 3.3 Groundwater resistomes from countries from different average income diverge

As a first variable, the average income of the country of sample origin was considered, as differing environmental/health standards, antibiotic usage patterns and sustainable agricultural practices between HICs and LMICs could affect the groundwater resistome. Based on the observed core ARG relative abundances, groundwater resistomes significantly differed between HICs and LMICs (Fig. 2A, PERMANOVA, R^2^=0.18, *p*=0.0001, n=29-39). Other sample origin-based factors, such as the depth of the groundwater samples did not affect the composition of resistomes (Table S3, PERMANOVA, R^2^=0.013, p=0.3, n=68) or in the case of population density in nearby regions had only a minor effect on ARG profiles (Table S3, PERMANOVA, R^2^=0.05, *p=*0.02, n=68). The core ARG classes displayed significantly higher relative abundance in groundwater metagenomes from LMICs when compared to HICs (Fig. 2B, Wilcoxon rank-sum test, *p*<0.05, n=29-39). Exclusively, β-lactam ARG relative abundance did not significantly differ between HICs and LMICs (Fig. 2B, Wilcoxon rank-sum test, *p>*0.05, n=29-39). At the individual ARG level a similar picture was observed: all core resistome ARGs were significantly higher relative abundant in groundwater metagenomes from LMICs compared to HICs (Fig. 2C, Wilcoxon-rank sum test, *p*<0.05, n=29-39), with the exception of the β-lactam ARG *bla*_OXA_ (Fig. 2C, Wilcoxon-rank sum test, *p*>0.05, n=29-39). Consequently, a clear difference in groundwater resistomes between HICs and LMICs with higher ARG relative abundances in LMICs was observed.

**Figure 2.**
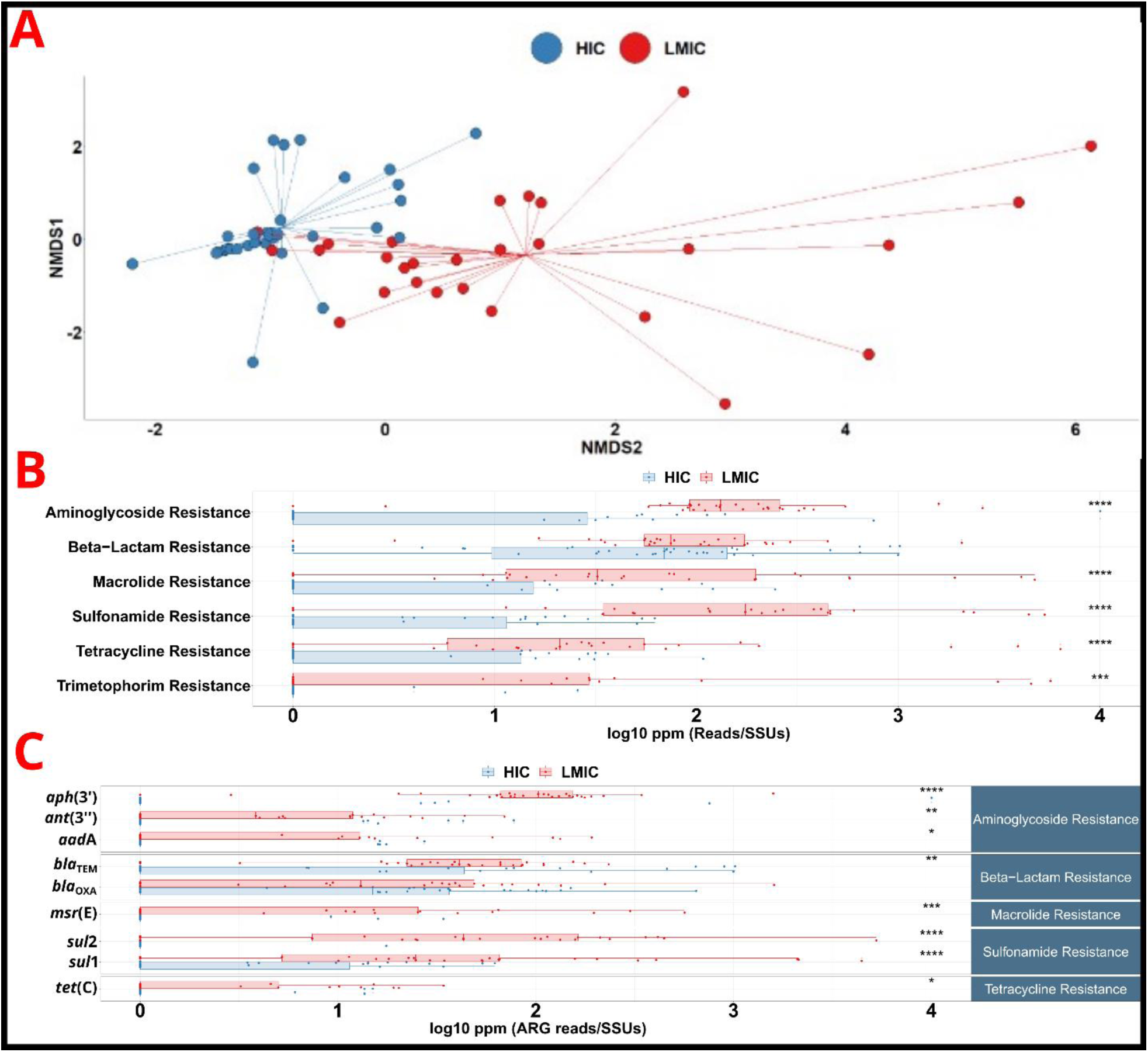
A) NMDS plot based on Euclidean Distance of the core resistome revealing differences in resistomes between high income countries (HIC) and lower-middle income countries (LMIC) (PERMANOVA, R^2^=0.18, *p*=0.0001, n=29-39). B) Core relative abundance at the ARGs summed at the antibiotic class level between HIC and LMIC countries (Wilcoxon rank-sum test, \**p*<0.05, \*\**p*<0.01, \*\*\**p*<0.001, \*\*\*\**p*<0.0001, n=29-39). Displayed are the summed relative abundances of all ARGs annotated based on their antibiotic classes, with a prevalence cut-off of appearance in at least 20% of samples. C) ARG abundance of core-ARGs between HIC and LMIC countries (Wilcoxon rank-sum test, *p*<0.05, n=29-39) with a prevalence cut-off of appearance in at least 20% of samples.

### 3.4 Higher abundances of mobile genetic element markers and core ARGs are correlated

As ARGs are often localized on MGEs, higher ARG relative abundance is regularly connected to an increased spread of MGEs in the respective bacterial communities. To evaluate if the dissemination of MGEs mirrors that of ARGs, we mapped the groundwater metagenomes to MGE databases and selected *int*I1*, kor*B, *tnp*A and general phage-abundance as MGE-markers that regularly link to ARGs. Similar to ARGs, MGE-markers exhibited significantly higher relative abundance in the groundwater from LMICs (Fig. 3A Wilcoxon rank-sum test, *p*<0.05, n=29-39). Out of the selected MGE markers, *tnp*A displayed the highest relative abundance, followed by phages, *kor*B and *int*I1. Furthermore, all core ARGs were significantly positively correlated with at least one MGE marker, while *sul*1 and *sul*2 correlated with all MGE markers (Fig. 3B).

**Figure 3.**
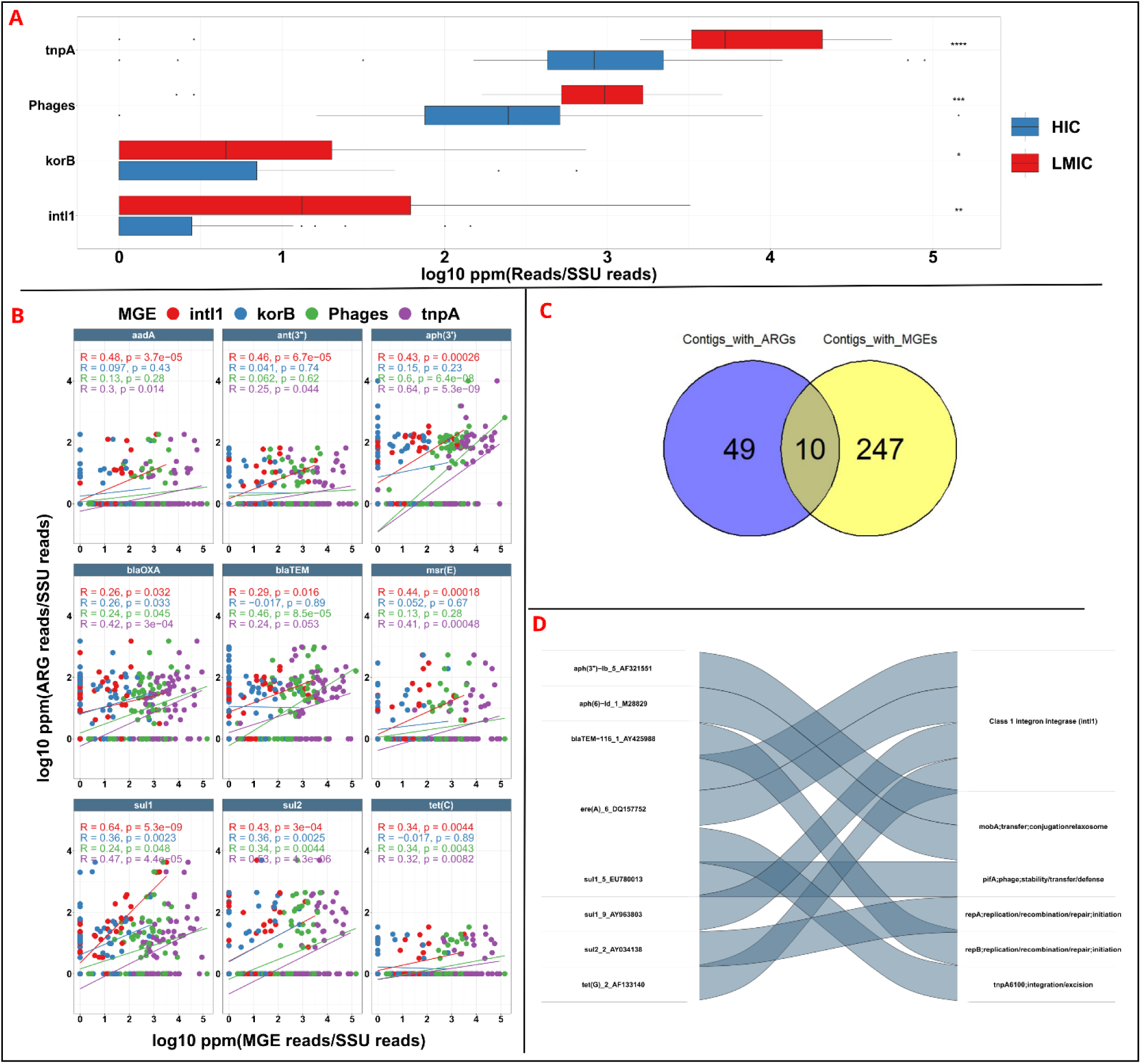
A) The relative abundance of the selected key MGE-markers in groundwater metagenomes from high income (HIC) and lower-middle income countries (LMIC) countries (Wilcoxon rank-sum test, \**p*<0.05, \*\**p*<0.01, \*\*\**p*<0.001, \*\*\*\**p*<0.0001, n=29-39). B) Spearman correlation and linear regression models of the relative abundance of core ARGs (Fig. 2C) and key MGEs (n=68). C) Contigs of groundwater metagenomes assembled via MEGAHIT. Following assembly, we perform alignment with DIAMOND to evaluated the occurrence of ARGs (via ResFinder) and MGEs (Mobile-OG) in the contigs. The overlap (number of different lines) indicates the number of contigs that contained both ARGs and MGE, and thus a potential co-localization. D) Sankey plot of ARGs and MGEs that were co-localized in the same contig. The numbers at the end and the letters, refer to the subtype of the gene as given by ResFinder database.

Based on this indication, we investigated if ARGs were indeed co-occurring on MGEs by assembling contigs from the groundwater metagenomes. In total, out of the 306 high-quality assembled contigs, 59 contained ARGs, while MGEs were present in 257 contigs (Fig. 3C). Ten contigs provided evidence for the localization of ARGs on MGEs (Fig. 3C). The gene *int*I1 was co-localized with ARGs *sul1* and erythromycin ARG *ere*(A) (Fig. 3D), despite *ere*(A) not having high enough prevalence across samples to be included in the core resistome. In addition, *int*I1 was also co-localized on a different contig with tetracycline resistance *tet*(G), a further ARG outside the core-resistome. Phage protein gene *pif*A was co-localized with *sul*1 (Fig. 3D), while *bla*_TEM_ and *sul*2 were co-localized with plasmid replication initiation protein genes *rep*A and *rep*B, indicating their conjugal transmission potential (Fig. 3D). In summary, higher MGE relative abundance was detected in groundwater from LMICs and a clear correlation between ARG and MGE relative abundance was established, with direct linkage of ARGs and MGEs evident based on contig-based analysis.

### 3.5 Bacterial community composition weakly associated with resistome composition

To evaluate if the difference in groundwater resistomes between HICs and LMICs could simply stem from an altered microbial community composition with naturally higher levels of ARGs, we performed SSU retrieval and classification via METAXA2. Similar to the resistome, bacterial community composition significantly differed between HICs and LMICs (Fig. 4A, PERMANOVA, average income: R^2^=0.12, *p*=0.0001, n=29-39). Furthermore, community composition and resistome composition were suggested to be linked based on a Mantel test on the dissimilarities of community composition and resistome composition (rho=0.17, *p=*0.02, Fig. 4B).

**Figure 4.**
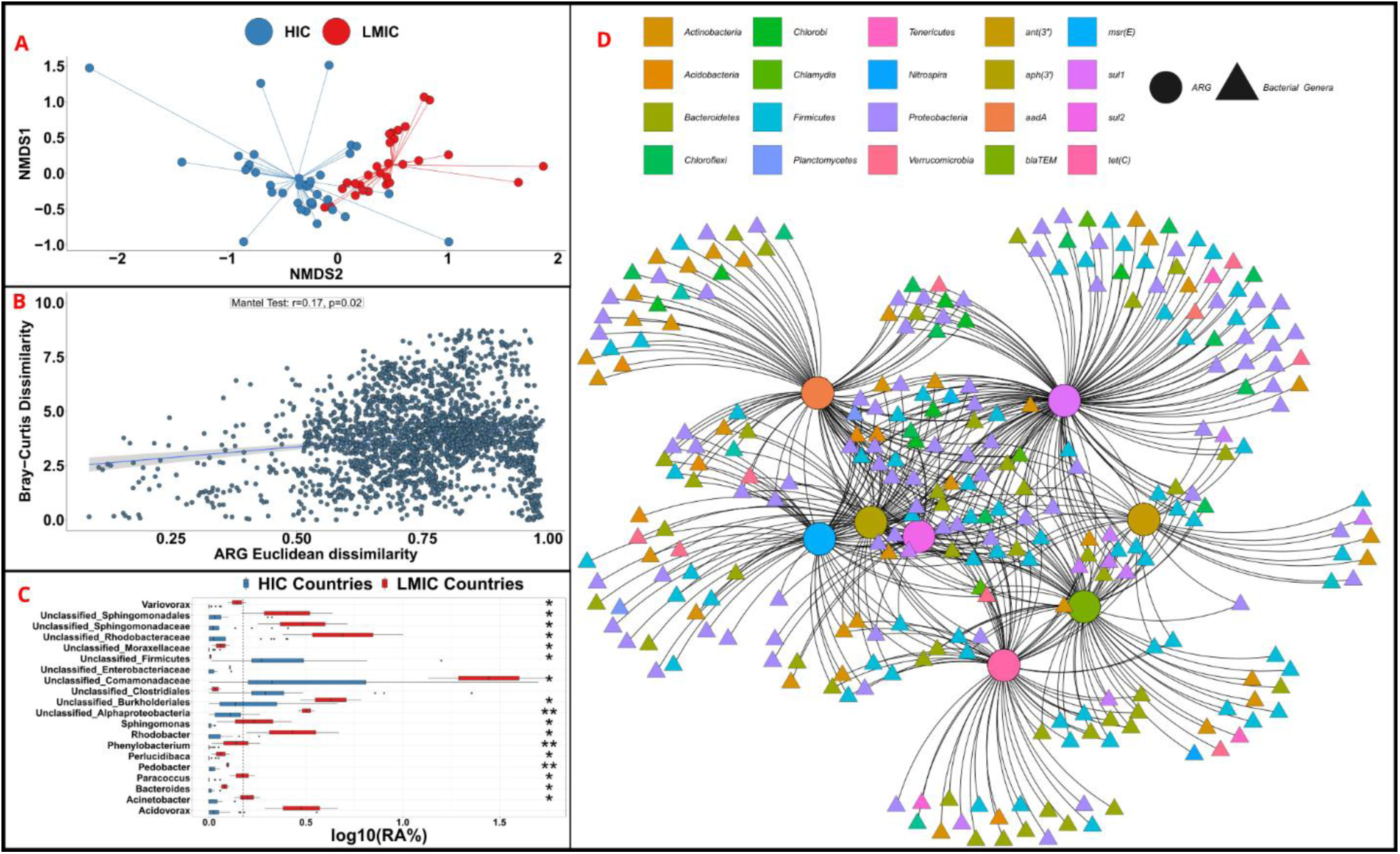
A) The bacterial community profile between HIC and LMIC based on Bray-Curtis distance (PERMANOVA, Average income: R^2^=0.12, *p*=0.0001, n=29-39). B) Mantel test with bacterial community and ARG compositions. C) The 20 most important bacterial taxa that differ between groundwater from HIC and LMIC. A random forest model was used to differentiate between groundwater metagenomes from HIC and LMIC in groundwater. Bacterial classification was performed at all taxonomic ranks from phylum until genus level and unclassified taxonomic ranks were summed at the latest unclassified level. The significance of differential abundance was assessed with Wilcoxon-rank sum test (\**p*<0.05, \*\**p*<0.01, \*\*\**p*<0.001). D) Correlation networks of core-ARGs with different bacterial genera based on Spearman correlation with Benjamini-Hochberg correction. The color legend shows the taxonomy of these genera at phylum level. Pairwise correlations were performed at a global dataset scale and only correlations with *p*<0.05 and *rho*>0 ARGs were subtracted for creating the network.

When investigating if individual ARGs were linked to specific bacterial taxa, integrated network analysis of bacterial genera and ARGs revealed that many core ARGs correlated with several bacterial genera from several phyla, including *Actinobacteria, Firmicutes* and *Proteobacteria*. The gene with the most correlated genera was *sul*1 (86 positive correlations in total), while *tet*(C) showed the lowest number of correlations (Fig. 4D).

However, when utilizing random forest algorithms to identify the most important taxa that contributed to the difference of groundwater metagenomes between HIC and LMIC (Fig. 4C), most of the genera having connections with ARGs in network analysis did not appear, except *Bacteroides* spp.. Among the bacterial genera that significantly differed based on Wilcoxon rank-sum tests (*p*<0.05, n=29-39) *Acinetobacter*, *Acidovorax, Variovorax, Bacteroides, Pedobacter, Sphingomonas, Perlucidibaca, Rhodobacter* and *Phenylobacterium* spp. were associated with LMICs. As our method, due to re-analysis of only short-read sequencing libraries, did not manage to classify all bacterial taxa at the genus level (Fig. S1), several unclassified bacterial taxa also significantly contributed to the observed differences (Fig. 4D, Wilcoxon rank-sum test, *p*<0.05, n=29-39). In conclusion, the bacterial community composition significantly differed between HICs and LMICs and this difference was correlated with the core ARGs, but no bacterial taxa highly accounting for the observed resistome differences could be identified in the co-occurrence network. Thus, the co-occurrence network along with the Mantel test indicated that bacterial community composition was only weakly associated with the resistome.

### 3.6 Relative abundance of natural producers of antimicrobials correlated with certain antibiotic classes

Aside from bacterial community composition, we aimed to further explore the underlying drivers of groundwater resistome diversity and relative abundance by evaluating whether ecological interactions with natural producers of antibiotics such as fungi and *Actinobacteria* could play a role in this rather pristine environment. We hence first evaluated the correlation of the core classes of ARGs with fungal relative abundance (i.e., fungal/bacterial SSUs in the metagenomes).

A clear correlation between fungal/bacterial relative abundance and beta-lactam resistance (Spearman rho=0.31, *p=*0.01, n=68, Fig. 5) was observed. The observed correlation was further verified using a linear mixed model: here, the original study from which the metagenomes were derived was set as a random effect variable to counter potential study-based biases (ARG Relative Abundance ∼ Fungal Relative Abundance + 1|Original_Study, *p*=0.044, n=68, Table S4). Moreover, the main bacterial antibiotic producers within the *Actinobacteria* phylum initially correlated exclusively with sulfonamide resistances and this correlation was negative (rho=-0.36, *p*=0.0024, n=68, Fig. 5). In contrast, when performing the linear mixed model accounting for study biases with ARG classes and *Actinobacteria*, *Actinobacteria* abundance significantly positively correlated with aminoglycosides, a class of antibiotics that many species classified as *Actinobacteria* produce (ARG Relative Abundance ∼ *Actinobacteria* Relative Abundance + 1|Original_Study, *p*=0.0366, n=68, Table S4) (Barka et al., 2016).

**Figure 5.**
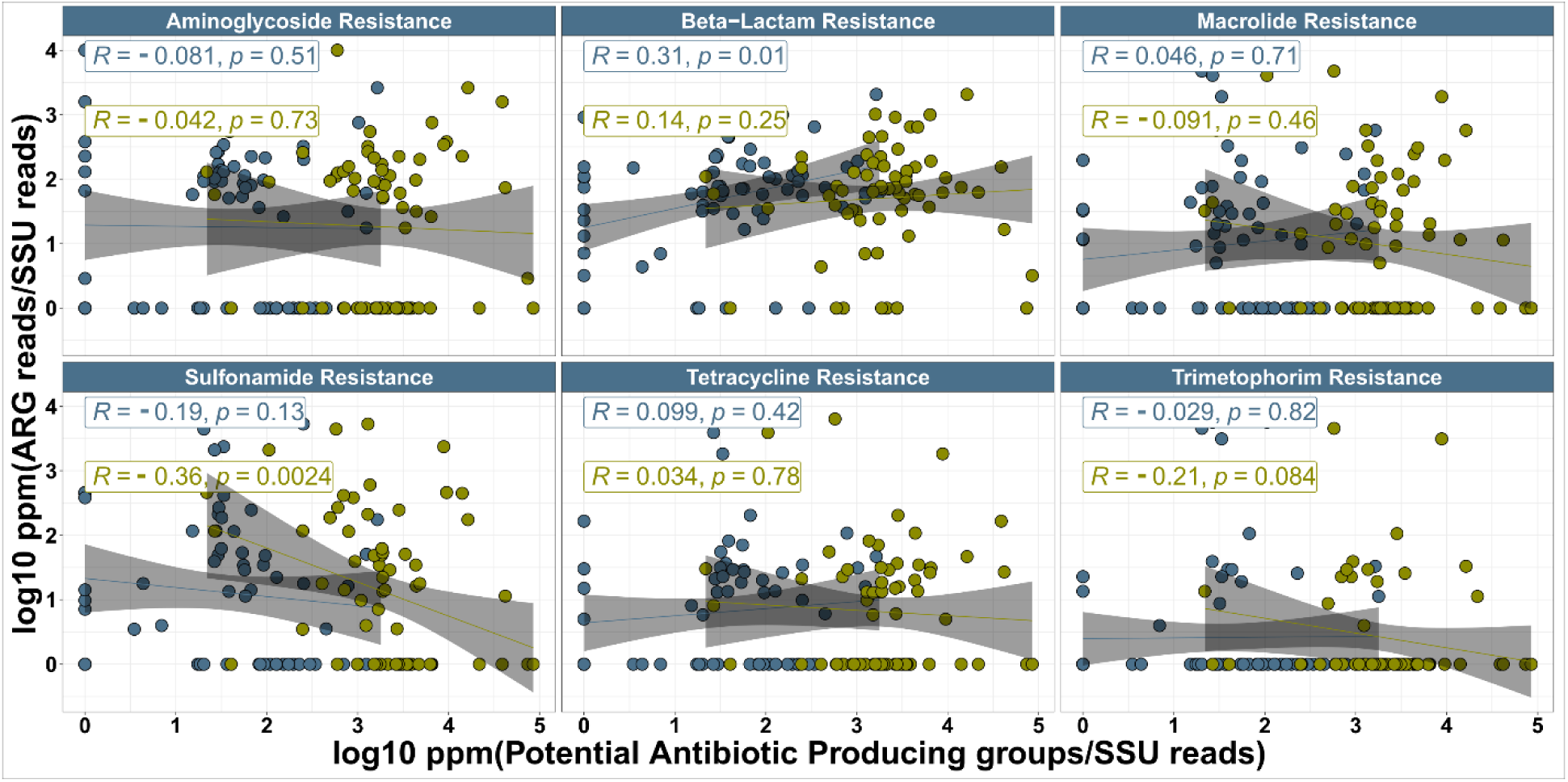
Correlations of potential antibiotic-producing groups (blue=Fungi, yellow=*Actinobacteria*) with ARG abundance at the antibiotic class level (Spearman correlation, n=68)

At the ARG level, none of the ARGs correlated with fungal or *Actinobacteria* abundance based on Spearman correlation (*p*>0.05, n=68, Fig. 5). Still, when using the mixed effect models to account for the study variances we detected significant positive correlations for the beta-lactam ARG *bla*_TEM_ and the aminoglycoside ARG *aad*A with both fungal and *Actinobacteria* abundance (ARG Relative Abundance ∼ *Actinobacteria* or Fungal Relative Abundance + 1|Original_Study, *p<*0.05, n=68, Table S4). In addition, two aminoglycoside ARG (*aph*(3’) and aadA) positively correlated with *Actinobacteria* abundance (ARG Relative Abundance ∼ *Actinobacteria* Relative Abundance + 1|Original_Study, *p*<0.05, n=68, Table S4). Consequently, the overall abundance of antibiotic-producing groups correlated with the levels of ARGs that confer resistance to β-lactam and aminoglycoside antibiotics, especially when we used models that take the individual study biases into account.

### 3.7 Anthropogenic factors explain the dissemination of antibiotic resistance with exception of β-lactam resistance which correlates with fungal and phage abundance

As outlined above, several different explanatory variables that show diverse correlations with resistome composition and ARG abundance were identified in individual analyses. However, to estimate the confounding effects of these different variables and the main drivers underlying resistome composition, we constructed a structural equation mixed effect model including all identified effectors and accounted for biases between the different studies. The model included ARG class abundance, MGEs, bacterial community composition, natural antibiotic producers, and average income of the sample origin country. The constructed structural equation model (RMSEA=0.28, Chi-square=37.16) confirmed that the average income most strongly explained the presence of sulfonamide, aminoglycoside, tetracycline and macrolide resistance genes (Fig. 6).

**Figure 6.**
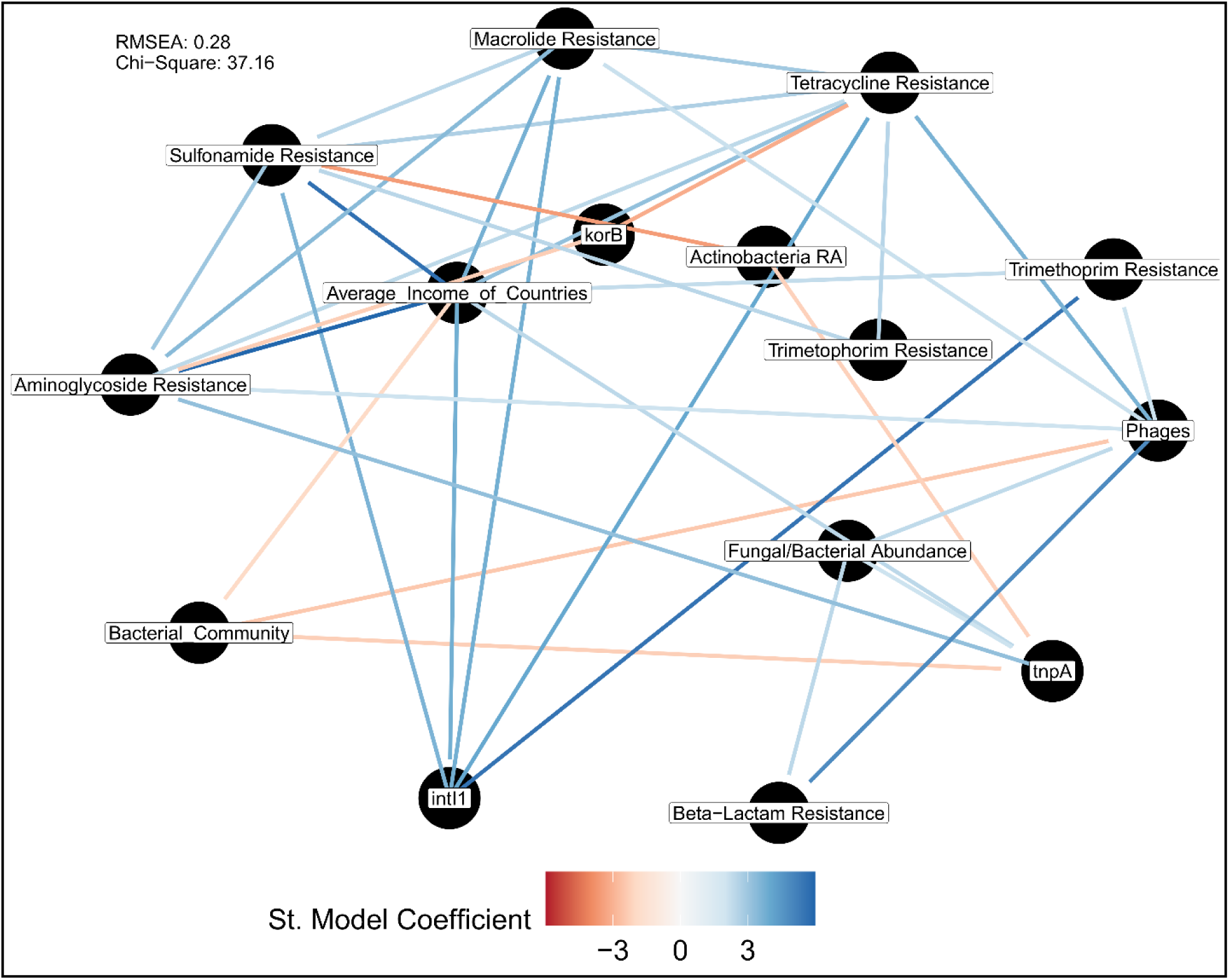
Structural equation mixed effect model based on partial least squares for evaluating the effect of multiple variables on the dissemination of the most frequent ARG classes. The model was fitted with log10-transformed and scaled abundances of reads as part per million for ARGs, MGEs and fungal/bacterial SSU. The model used the original study as a random variable to account for the biases between the different studies. The estimate of model co-efficient (α) indicates positive or negative correlations based on linear regression with unscaled coefficients. For evaluating the compositional changes of bacterial community in the model, we used the first principal component from a fitted PCoA based on Bray-Curtis distance. Only significant positive (blue) or negative (red) relationships with *p*<0.05 are displayed (n=68). Model shows standardized regression coefficient based on linear regression.

Unsurprisingly, the average income was also associated with the abundance of *intI1*, a common indicator gene for anthropogenic influence on environmental samples (Gillings, 2017) (Fig. 6). It is worth noting that many antibiotic classes exhibited correlations among each other, such as aminoglycoside, tetracycline and macrolide resistances, confirming that global abiotic factors affecting the entire resistome such as the average income, were more important in shaping the groundwater resistome than biotic interactions working on the individual gene level.

However, β-lactam resistance was mainly explained by biotic interactions with fungal relative abundance. Meanwhile, *Actinobacteria*, main bacterial producers of antimicrobials, had a slightly negative correlation with sulfonamide resistance, but still displayed no positive correlation with any antibiotic resistance class (Fig. 6). Notably, while community composition strongly depended on the average income, it did not directly affect the abundance of any ARG class (Fig. 6). This indicates the low explanatory power of the microbiome on the resistome composition that might be lost when confounding, deterministic variables and processes are present. At the MGE level, sulfonamide resistance was partly explained by *intI1* abundance (Fig. 6), while, the presence of phages was positively correlated with β-lactam, aminoglycoside, tetracycline, macrolide and trimethoprim resistance (Fig. 6). The co-localization trends predicted by the structural equation model, for sulfonamide resistance with *intI1* were previously confirmed through contig assembly, displaying the reliability of this approach (Fig. 5D & 6). However, certain ARG-MGE connections identified in the herein assembled contigs (e.g., *sul1* co-localization with phages) seemed not to significantly affect the ARG’s relative abundance on the global scale.

## 4. Discussion

Groundwater environments constitute an important drinking water resource, hence gaining insights into their microbial composition is critical for increasing sustainability of water resources. Herein, we disentangle the combined effect of anthropogenic, biotic, and environmental drivers on the AMR distribution in groundwater environments by re-analyzing already existing metagenomes. Using this *in silico* approach, we demonstrated that anthropogenic factors such as the average income of the sample’s country of origin play a major role and affect the entire resistome. Thus, mitigation policies on antibiotic use, which are more prevalent in HICs, seem to be reflected in decreased AMR abundances in groundwater environments. In addition, biotic factors such as interactions with fungi or phages contributed to the dissemination of different individual antibiotic classes or individual ARGs and ARG-associated MGEs in the global resistome, such as demonstrated for β-lactam resistance. Hence, our results indicate that a complex interplay of anthropogenic activities and biotic interactions shape the global groundwater resistome.

Severe differences in groundwater resistomes between HICs and LMICs were observed, with LMIC countries displaying higher levels of resistance. HICs have regularly adopted stricter regulations regarding antibiotic use, which translates into lower rates of antibiotic consumption for patients, less antibiotic use for livestock production, higher standards for waste disposal from antibiotic manufacturing factories, and more efficient sanitation standards including wastewater infrastructure (Theuretzbacher et al., 2017). Since already the infiltration of only low concentrations of selective compounds has the potential to severely affect groundwater microbiomes (Kampouris et al., 2022), different practices in livestock production and waste disposal (Heuer et al., 2011; Heuer and Smalla, 2007; Jechalke et al., 2016) or increased use of groundwater aquifer recharge through treated wastewater, that is more regularly used in LMIC countries (Zhang et al., 2020), might have contributed to the increased AMR levels observed in those countries. While the presence and abundance of ARGs in different environments can regularly be explained by the degree of direct fecal contamination (Karkman et al., 2019), here no crAssphage reads were detected in any of the studied metagenomes, indicating that the infiltration of fecal organisms is, if at all, of very minor relevance for the rather pristine groundwater environments and can be excluded as an explanatory variable. Still the potentially increased anthropogenic influence is mirrored in the detection of elevated levels of the Class1 integron *int*I1 gene in LMIC groundwater metagenomes. *int*I1 has regularly been suggested as an indicator of anthropogenic impact (Gillings et al., 2015; Smalla et al., 2018) and was here also positively correlated with several types of ARGs, especially sulfonamide ARG *sul*1 which is regularly encoded as part of Class1 integron cassettes (De La Cruz Barron et al., 2023). Furthermore, co-localization of other ARGs with *intI*1 (i.e., *ere*(A) and *tet*(G)) was here confirmed based on assembly analysis displaying the potential of co-selection and co-spread of multiple ARGs.

Theoretically, the assembly of metagenomes enables the co-localization of ARGs with MGEs, but many connections are often overlooked due to sequencing depth limitations (Abramova et al., 2023). These limitations were prominent in the global groundwater metagenomes, where only ten contigs had ARGs and MGEs co-located. This observed co-localization of ARGs and MGEs is indicative of an increased future risk (Bengtsson-Palme et al., 2023; De La Cruz Barron et al., 2023; Zhang et al., 2021). Using culture-based methods for the exogenous capturing of ARG-containing plasmids (Blau et al., 2018; Shintani, 2023), might lead to more qualitative information regarding which MGEs are particularly associated with the spread of ARGs in groundwater compared to the here-used metagenomic reanalysis and assembly based approach. When assessing if biotic interactions could also be shaping the groundwater resistomes, we detected antibiotic class-specific positive correlations of ARGs with those microbial groups hosting the majority of antibiotic producers, such as fungi or *Actinobacteria* (Aly et al., 2011; Barka et al., 2016). Based on the structural equation model, a clear correlation of fungal abundance with β-lactam resistance, the only class of antibiotics that was not explained based on the average income, was detected. Since fungi can produce β-lactams (Aly et al., 2011), this relationship might originate from competition between fungi and bacteria, or quorum-sensing-based interactions (Romero et al., 2011). Interestingly, fungal abundance and β-lactam resistance correlated also with phage abundance. Previous research has indicated that fungal hyphae can assist in the transfer of phages (Ghanem et al., 2019) and increase their infection rates, which could be connected to these correlations. Contrary to fungi, *Actinobacteria*, the main bacterial group of antibiotic producers (Barka et al., 2016), did finally not play any significant role in shaping the global groundwater resistomes in the structural equation model, although initially weak correlations with aminoglycoside resistance were indicated using simpler correlation-based approaches.

Finally, groundwater is characterized by the absence of heavy selection pressures favoring selection for resistant bacteria. It has previously been reported that in rather pristine, low-selection environments such as drinking water reservoirs bacterial community and resistome composition are regularly unlinked (Fang et al., 2019). While both network analysis and the Mantel test indicated a certain linkage between the two, the structural equation model revealed that bacterial community composition indeed had a low explanatory power on the groundwater resistome composition. Still, the difference based on average income similar to the resistome strongly affected the β-diversity. This raises questions about whether other microbial risk factors such as the occurrence of pathogens might be favored (Ferrer et al., 2020) in either type of groundwater microbiome. Here, *Acinetobacter* spp., a genus that includes several pathogenic and multidrug resistance strains (Hernández-González and Castillo-Ramírez, 2020; Kyriakidis et al., 2021), displayed a far increased abundance in LMICs. Apart from their clinical relevance *Acinetobacter* spp., a group of obligate aerobes (Jung and Park, 2015), are widespread in soil (Adegoke et al., 2012) and as indicated here in groundwater environments. Since *Acinetobacter* spp. has regularly been found to degrade pollutants (Dahal et al., 2023), this increase might reflect the overall higher level of organic pollutants in LMICs. This is also supported by the higher abundance of *Sphingomonas* spp. in LMICs, a genus that also includes strains with a high capacity for pollutant degradation (Asaf et al., 2020). Metagenomics based on short-read technologies might have high limitations on verifying if the observed strains from these genera do indeed have pathogenic potential (Tedersoo et al., 2021). Still, the wide occurrence of genera that include several potential pathogenic strains should not be neglected.

## 5. Conclusions

In summary, we demonstrate that the re-analysis of publicly available environmental microbiome data is a valuable source for testing and creating general ecological hypotheses and elucidating potential global factors shaping microbiomes and resistomes. We here applied it to the relatively pristine groundwater microbiome to determine how different environmental conditions, anthropogenic factors, and biotic relationships affect the global groundwater resistome. Specifically, we demonstrate that the average income of the country of sample origin significantly shapes the resistome composition. Meanwhile, we demonstrate that the abundance of certain groups of ARGs can be predicted based on microbe-microbe interactions. Specifically, we demonstrate that the bacterial/fungal SSU ratio could act as an indicator of the abundance of β-lactam resistance in groundwater environments. We expect that with the increase of publicly available metagenomic data, such *in silico* meta-analyses will be able to further identify ecological interactions in understudied environments in the future, and can be used for creating hypotheses that can be tested later in lab and field experiments. Furthermore, the results from this study contribute to a necessary baseline for future groundwater AMR surveillance.

## 6. Conflict of interest

The authors declare no conflict of interest.

## 7. Acknowledgements

We deeply thank the researchers who uploaded and provided their sequencing data and metadata in the NCBI SRA databases, as without their contribution this study would have been impossible.

## 8. Funding

This work was supported by the JPI AMR (EMBARK and SEARCHER; JPIAMR2019-109 and JPIAMR2023-DISTOMOS-016) funded by the Bundesministerium für Bildung und Forschung under grant numbers F01KI1909A & 01KI2404A and the Swedish Research Council (VR; grants 2019-00299 and 2023-01721). TUB, BL & UK acknowledge funding from the German-Chinese collaboration project Explore-AMR funded by the Bundesministerium für Bildung und Forschung under grant number 01DO2200 and the National Key R&D Program of China under grant number 2022YFE0103200. JBP acknowledges funding under the frame of the Data-Driven Life Science (DDLS) program supported by the Knut and Alice Wallenberg Foundation (KAW 2020.0239) and the Swedish Foundation for Strategic Research (FFL21-0174). Responsibility for the information and views expressed in the manuscript lies entirely with the authors.

## 9. Data availability

Data and the R and shell scripts for the workflow of analysis have been uploaded to GitHub under https://github.com/JonKampouris/groundwater_Resistome. A list of the accession numbers of the used groundwater metagenomes can be found in the supplementary materials.

## Notes

https://github.com/JonKampouris/GW_Resistome

### Competing Interest Statement

The authors have declared no competing interest.

### Summary of Updates

Analysis was extended to include more metagenomes and additional metadata. Authors have been added.

## 10. References

Abramova, A., Karkman, A., Bengtsson-Palme, J., 2023. Metagenomic assemblies tend to break around antibiotic resistance genes. bioRxiv 2023.12.13.571436. 10.1101/2023.12.13.571436

Adegoke, A.A., Mvuyo, T., Okoh, A.I., 2012. Ubiquitous Acinetobacter species as beneficial commensals but gradually being emboldened with antibiotic resistance genes. J. Basic Microbiol. 52, 620–627. 10.1002/JOBM.201100323

Aly, A.H., Debbab, A., Proksch, P., 2011. Fungal endophytes: Unique plant inhabitants with great promises. Appl. Microbiol. Biotechnol. 90, 1829–1845. 10.1007/s00253-011-3270-y

Asaf, S., Numan, M., Khan, A.L., Al-Harrasi, A., 2020. Sphingomonas: from diversity and genomics to functional role in environmental remediation and plant growth. Crit. Rev. Biotechnol. 40, 138–152. 10.1080/07388551.2019.1709793

Bahram, M., Hildebrand, F., Forslund, S.K., Anderson, J.L., Soudzilovskaia, N.A., Bodegom, P.M., Bengtsson-Palme, J., Anslan, S., Coelho, L.P., Harend, H., Huerta-Cepas, J., Medema, M.H., Maltz, M.R., Mundra, S., Olsson, P.A., Pent, M., Põlme, S., Sunagawa, S., Ryberg, M., Tedersoo, L., Bork, P., 2018. Structure and function of the global topsoil microbiome. Nature 560, 233–237. 10.1038/s41586-018-0386-6

Barka, E.A., Vatsa, P., Sanchez, L., Gaveau-vaillant, N., Jacquard, C., Klenk, H., Clément, C., Ouhdouch, Y., Wezel, P. Van, 2016. Taxonomy, Physiology, and Natural Products of Actinobacteria 21. 10.1128/MMBR.00019-15.Address

Bates, D., Mächler, M., Bolker, B.M., Walker, S.C., 2015. Fitting linear mixed-effects models using lme4. J. Stat. Softw. 67. 10.18637/jss.v067.i01

Bengtsson-Palme, J., Abramova, A., Berendonk, T.U., Coelho, L.P., Forslund, S.K., Gschwind, R., Heikinheimo, A., Jarquín-Díaz, V.H., Khan, A.A., Klümper, U., Löber, U., Nekoro, M., Osińska, A.D., Ugarcina Perovic, S., Pitkänen, T., Rødland, E.K., Ruppé, E., Wasteson, Y., Wester, A.L., Zahra, R., 2023. Towards monitoring of antimicrobial resistance in the environment: For what reasons, how to implement it, and what are the data needs? Environ. Int. 178. 10.1016/j.envint.2023.108089

Bengtsson-Palme, J., Hartmann, M., Eriksson, K.M., Pal, C., Thorell, K., Larsson, D.G.J., Nilsson, R.H., 2015. metaxa2: Improved identification and taxonomic classification of small and large subunit rRNA in metagenomic data. Mol. Ecol. Resour. 15, 1403–1414. 10.1111/1755-0998.12399

Blau, K., Bettermann, A., Jechalke, S., Fornefeld, E., Vanrobaeys, Y., Stalder, T., Top, E., Smalla, K., 2018. The transferable resistome of produce. bioRxiv 9, 1–15. 10.1101/350629

Bortolaia, V., Kaas, R.S., Ruppe, E., Roberts, M.C., Schwarz, S., Cattoir, V., Philippon, A., Allesoe, R.L., Rebelo, A.R., Florensa, A.F., Fagelhauer, L., Chakraborty, T., Neumann, B., Werner, G., Bender, J.K., Stingl, K., Nguyen, M., Coppens, J., Xavier, B.B., Malhotra-Kumar, S., Westh, H., Pinholt, M., Anjum, M.F., Duggett, N.A., Kempf, I., Nykäsenoja, S., Olkkola, S., Wieczorek, K., Amaro, A., Clemente, L., Mossong, J., Losch, S., Ragimbeau, C., Lund, O., Aarestrup, F.M., 2020. ResFinder 4.0 for predictions of phenotypes from genotypes. J. Antimicrob. Chemother. 75, 3491–3500. 10.1093/jac/dkaa345

Brown-Jaque, M., Calero-Cáceres, W., Muniesa, M., 2015. Transfer of antibiotic-resistance genes via phage-related mobile elements. Plasmid 79, 1–7. 10.1016/J.PLASMID.2015.01.001

Brown, C.L., Mullet, J., Hindi, F., Stoll, J.E., Gupta, S., Choi, M., Keenum, I., Vikesland, P., Pruden, A., Zhang, L., 2022. mobileOG-db: a Manually Curated Database of Protein Families Mediating the Life Cycle of Bacterial Mobile Genetic Elements. Appl. Environ. Microbiol. 88. 10.1128/AEM.00991-22/SUPPL_FILE/AEM.00991-22-S0004.XLSX

Buchfink, B., Xie, C., Huson, D.H., 2014. Fast and sensitive protein alignment using DIAMOND. Nat. Methods 12, 59–60. 10.1038/nmeth.3176

Cock, P.J.A., Antao, T., Chang, J.T., Chapman, B.A., Cox, C.J., Dalke, A., Friedberg, I., Hamelryck, T., Kauff, F., Wilczynski, B., De Hoon, M.J.L., 2009. Biopython: Freely available Python tools for computational molecular biology and bioinformatics. Bioinformatics 25, 1422–1423. 10.1093/bioinformatics/btp163

Coelho, L.P., Alves, R., Monteiro, P., Huerta-Cepas, J., Freitas, A.T., Bork, P., 2019. NG-meta-profiler: Fast processing of metagenomes using NGLess, a domain-specific language. Microbiome 7, 1–10. 10.1186/s40168-019-0684-8

Csárdi, G., Nepusz, T., Müller, K., Horvát, S., Traag, V., Zanini, F., Noom, D., 2023. igraph for R: R interface of the igraph library for graph theory and network analysis. Zenodo. Available online https://CRAN.R-project.org/package=igraph (accessed March 30, 2023). 10.5281/ZENODO.10681749

Dahal, U., Paul, K., Gupta, S., 2023. The multifaceted genus Acinetobacter: from infection to bioremediation. J. Appl. Microbiol. 134, 1–18. 10.1093/JAMBIO/LXAD145

David Sjoberg, 2023. davidsjoberg/ggsankey: Make sankey, alluvial and sankey bump plots in ggplot [WWW Document]. URL https://github.com/davidsjoberg/ggsankey (accessed 6.7.24).

De La Cruz Barron, M., Kneis, D., Elena, A.X., Bagra, K., Berendonk, T.U., Klümper, U., 2023. Quantification of the mobility potential of antibiotic resistance genes through multiplexed ddPCR linkage analysis. FEMS Microbiol. Ecol. 99, 1–10. 10.1093/femsec/fiad031

Fang, P., Peng, F., Gao, X., Xiao, P., Yang, J., 2019. Decoupling the Dynamics of Bacterial Taxonomy and Antibiotic Resistance Function in a Subtropical Urban Reservoir as Revealed by High-Frequency Sampling 10, 1–12. 10.3389/fmicb.2019.01448

Ferrer, N., Folch, A., Masó, G., Sanchez, S., Sanchez-vila, X., 2020. What are the main factors in fl uencing the presence of faecal bacteria pollution in groundwater systems in developing countries ? J. Contam. Hydrol. 228, 103556. 10.1016/j.jconhyd.2019.103556

Flynn, T.M., Flynn, T.M., Sanford, R.A., Ryu, H., Bethke, C.M., Levine, A.D., 2013. Functional microbial diversity explains groundwater chemistry in a pristine aquifer Functional microbial diversity explains groundwater chemistry in a pristine aquifer. BMC Microbiol. 13, 1.

François Briatte, Michał Bojanowski, Mickaël Canouil, Zachary Charlop-Powers, Jacob C. Fisher, Kipp Johnson, Tyler Rinker, 2024. Geometries to Plot Networks with “ggplot2” [R package ggnetwork version 0.5.13].

Gatica, J., Yang, K., Pagaling, E., Jurkevitch, E., Yan, T., Cytryn, E., 2015. Resistance of undisturbed soil microbiomes to ceftriaxone indicates extended spectrum β-lactamase activity. Front. Microbiol. 6, 1–11. 10.3389/fmicb.2015.01233

Ghanem, N., Stanley, C.E., Harms, H., Chatzinotas, A., Wick, L.Y., 2019. Mycelial E ff ects on Phage Retention during Transport in a Micro fl uidic Platform. 10.1021/acs.est.9b03502

Gillings, M.R., 2017. Class 1 integrons as invasive species. Curr. Opin. Microbiol. 38, 10–15. 10.1016/j.mib.2017.03.002

Gillings, M.R., Gaze, W.H., Pruden, A., Smalla, K., Tiedje, J.M., Zhu, Y.G., 2015. Using the class 1 integron-integrase gene as a proxy for anthropogenic pollution. ISME J. 9, 1269–1279. 10.1038/ismej.2014.226

Griebler, C., Avramov, M., 2015. Groundwater ecosystem services : a review 34, 355–367. 10.1086/679903.

Griebler, C., Lueders, T., 2009. Microbial biodiversity in groundwater ecosystems. Freshw. Biol. 54, 649–677. 10.1111/j.1365-2427.2008.02013.x

Hadley Wickham, 2016. ggplot2: Elegant Graphics for Data Analysis - Hadley Wickham - Βιβλία Google.

Hernández-González, I.L., Castillo-Ramírez, S., 2020. Antibiotic-resistant Acinetobacter baumannii is a One Health problem. The Lancet Microbe 1, e279. 10.1016/S2666-5247(20)30167-1

Hernando-Amado, S., Coque, T.M., Baquero, F., Martínez, J.L., 2019. Defining and combating antibiotic resistance from One Health and Global Health perspectives. Nat. Microbiol. 4, 1432–1442. 10.1038/s41564-019-0503-9

Herrmann, M., Wegner, C.E., Taubert, M., Geesink, P., Lehmann, K., Yan, L., Lehmann, R., Totsche, K.U., Küsel, K., 2019. Predominance of Cand. Patescibacteria in groundwater is caused by their preferential mobilization from soils and flourishing under oligotrophic conditions. Front. Microbiol. 10, 1–15. 10.3389/fmicb.2019.01407

Heuer, H., Schmitt, H., Smalla, K., 2011. Antibiotic resistance gene spread due to manure application on agricultural fields. Curr. Opin. Microbiol. 14, 236–243. 10.1016/j.mib.2011.04.009

Heuer, H., Smalla, K., 2007. Manure and sulfadiazine synergistically increased bacterial antibiotic resistance in soil over at least two months. Environ. Microbiol. 9, 657–666. 10.1111/j.1462-2920.2006.01185.x

Hosain, M.Z., Lutful Kabir, S.M., Kamal, M.M., 2021. Antimicrobial uses for livestock production in developing countries. Vet. World 14, 210. 10.14202/VETWORLD.2021.210-221

Jechalke, S., Radl, V., Schloter, M., Heuer, H., Smalla, K., 2016. Do drying and rewetting cycles modulate effects of sulfadiazine spiked manure in soil? FEMS Microbiol. Ecol. 92, 1–7. 10.1093/femsec/fiw066

Johnston, S.R., Boddy, L., Weightman, A.J., 2016. Bacteria in decomposing wood and their interactions with wood-decay fungi. FEMS Microbiol. Ecol. 92, 1–12. 10.1093/femsec/fiw179

Jung, J., Park, W., 2015. Acinetobacter species as model microorganisms in environmental microbiology: current state and perspectives. Appl. Microbiol. Biotechnol. 2015 996 99, 2533–2548. 10.1007/S00253-015-6439-Y

Kampouris, I.D., Alygizakis, N., Klümper, U., Agrawal, S., Lackner, S., Cacace, D., Kunze, S., Thomaidis, N.S., Slobdonik, J., Berendonk, T.U., 2022. Elevated levels of antibiotic resistance in groundwater during treated wastewater irrigation associated with infiltration and accumulation of antibiotic residues. J. Hazard. Mater. 423, 127155. 10.1016/j.jhazmat.2021.127155

Karkman, A., Pärnänen, K., Larsson, D.G.J., 2019. Fecal pollution can explain antibiotic resistance gene abundances in anthropogenically impacted environments. Nat. Commun. 10, 1–8. 10.1038/s41467-018-07992-3

Kyriakidis, I., Vasileiou, E., Pana, Z.D., Tragiannidis, A., 2021. Acinetobacter baumannii Antibiotic Resistance Mechanisms. Pathog. 2021, Vol. 10, Page 373 10, 373. 10.3390/PATHOGENS10030373

Laxminarayan, R., Duse, A., Wattal, C., Zaidi, A.K.M., Wertheim, H.F.L., Sumpradit, N., Vlieghe, E., Hara, G.L., Gould, I.M., Goossens, H., Greko, C., So, A.D., Bigdeli, M., Tomson, G., Woodhouse, W., Ombaka, E., Peralta, A.Q., Qamar, F.N., Mir, F., Kariuki, S., Bhutta, Z.A., Coates, A., Bergstrom, R., Wright, G.D., Brown, E.D., Cars, O., 2013. Antibiotic resistance-the need for global solutions. Lancet Infect. Dis. 13, 1057–1098. 10.1016/S1473-3099(13)70318-9

Li, D., Liu, C.M., Luo, R., Sadakane, K., Lam, T.W., 2015. MEGAHIT: An ultra-fast single-node solution for large and complex metagenomics assembly via succinct de Bruijn graph. Bioinformatics 31, 1674–1676. 10.1093/BIOINFORMATICS/BTV033

Li, H., 2013. Aligning sequence reads, clone sequences and assembly contigs with BWA-MEM 00, 1–3.

Li, H., Durbin, R., 2010. Fast and accurate long-read alignment with Burrows-Wheeler transform. Bioinformatics 26, 589–595. 10.1093/bioinformatics/btp698

Linares, J.F., Gustafsson, I., Baquero, F., Martinez, J.L., 2006. Antibiotics as intermicrobiol signaling agents instead of weapons. Proc. Natl. Acad. Sci. U. S. A. 103, 19484–19489. 10.1073/pnas.0608949103

Linlin Yan, 2023. Package “ggvenn”:An easy-to-use way to draw pretty venn diagram by “ggplot2”.

Martin, M., 2011. Cutadapt removes adapter sequences from high-throughput sequencing reads. EMBnet.journal 17, 10–12.

Nawaz, A., Purahong, W., Lehmann, R., Herrmann, M., Totsche, K.U., Küsel, K., Wubet, T., Buscot, F., 2018. First insights into the living groundwater mycobiome of the terrestrial biogeosphere. Water Res. 145, 50–61. 10.1016/j.watres.2018.07.067

Oksanen J, Simpson G, Blanchet F, Kindt R, Legendre P, Minchin P, O’Hara R, Solymos P, Stevens M, Szoecs E, Wagner H, Barbour M, Bedward M, Bolker B, Borcard D, Carvalho G, Chirico M, De Caceres M, Durand S, E.H., Fitz John R, Friendly M, Furneaux B, Hannigan G, Hill M, Lahti L, McGlinn D, Ouellette M, Ribeiro Cunha E, S.T., Stier A., Ter Braak C., W.J.., 2022. vegan: Community Ecology Package [WWW Document].

Pärnänen, K., Karkman, A., Hultman, J., Lyra, C., Bengtsson-Palme, J., Larsson, D.G.J., Rautava, S., Isolauri, E., Salminen, S., Kumar, H., Satokari, R., Virta, M., 2018. Maternal gut and breast milk microbiota affect infant gut antibiotic resistome and mobile genetic elements. Nat. Commun. 2018 91 9, 1–11. 10.1038/s41467-018-06393-w

R Core Team, 2021. R: A language and environment for statistical computing. [WWW Document]. URL https://www.r-project.org/ (accessed 6.7.24).

Retter, A., Karwautz, C., Griebler, C., 2021. Groundwater microbial communities in times of climate change. Curr. Issues Mol. Biol. 41, 509–538. 10.21775/cimb.041.509

Romero, D., Traxler, M.F., Daniel, L., Kolter, R., 2011. Antibiotics as Signal Molecules 5492– 5505.

Rosseel, Y., 2012. lavaan: An R Package for Structural Equation Modeling. J. Stat. Softw. 48, 1–36. 10.18637/JSS.V048.I02

Schlüter, A., Szczepanowski, R., Pühler, A., Top, E.M., 2007. Genomics of IncP-1 antibiotic resistance plasmids isolated from wastewater treatment plants provides evidence for a widely accessible drug resistance gene pool. FEMS Microbiol. Rev. 31, 449–477. 10.1111/J.1574-6976.2007.00074.X

Shintani, M., 2023. Integrons, transposons and IS elements promote diversification of multidrug resistance plasmids and adaptation of their hosts to antibiotic pollutants from pharmaceutical companies Antibiotic abbreviations 3035–3051. 10.1111/1462-2920.16481

Smalla, K., Cook, K., Djordjevic, S.P., Klümper, U., Gillings, M., 2018. Environmental dimensions of antibiotic resistance: assessment of basic science gaps. FEMS Microbiol. Ecol. 94, 1–6. 10.1093/femsec/fiy195

Sommermann, L., Geistlinger, J., Wibberg, D., Deubel, A., Zwanzig, J., Babin, D., Schlüter, A., Schellenberg, I., 2018. Fungal community profiles in agricultural soils of a long-term field trial under different tillage, fertilization and crop rotation conditions analyzed by high-throughput ITS-amplicon sequencing. PLoS One 13, e0195345. 10.1371/JOURNAL.PONE.0195345

Sonthiphand, P., Ruangroengkulrith, S., Mhuantong, W., Charoensawan, V., Chotpantarat, S., Boonkaewwan, S., 2019. Metagenomic insights into microbial diversity in a groundwater basin impacted by a variety of anthropogenic activities. Environ. Sci. Pollut. Res. 26, 26765–26781. 10.1007/s11356-019-05905-5

Szekeres, E., Chiriac, C.M., Baricz, A., Szőke-Nagy, T., Lung, I., Soran, M.L., Rudi, K., Dragos, N., Coman, C., 2018. Investigating antibiotics, antibiotic resistance genes, and microbial contaminants in groundwater in relation to the proximity of urban areas. Environ. Pollut. 236, 734–744. 10.1016/j.envpol.2018.01.107

Tedersoo, L., Albertsen, M., Anslan, S., 2021. Perspectives and Bene fi ts of High-Throughput Long-Read Sequencing in Microbial Ecology. Appl. Environ. Microbiol. 87. 10.1128/AEM.00626-21.

Theuretzbacher, U., Årdal, C., Harbarth, S., 2017. Linking sustainable use policies to novel economic incentives to stimulate antibiotic research and development. Infect. Dis. Rep. 9, 28–31. 10.4081/idr.2017.6836

Vaz-Moreira, I., Nunes, O.C., Manaia, C.M., 2014. Bacterial diversity and antibiotic resistance in water habitats: Searching the links with the human microbiome. FEMS Microbiol. Rev. 38, 761–778. 10.1111/1574-6976.12062

Wickham, H., Averick, M., Bryan, J., Chang, W., McGowan, L., François, R., Grolemund, G., Hayes, A., Henry, L., Hester, J., Kuhn, M., Pedersen, T., Miller, E., Bache, S., Müller, K., Ooms, J., Robinson, D., Seidel, D., Spinu, V., Takahashi, K., Vaughan, D., Wilke, C., Woo, K., Yutani, H., 2019. Welcome to the Tidyverse. J. Open Source Softw. 4, 1686. 10.21105/joss.01686

Zaouri, N., Jumat, M.R., Cheema, T., Hong, P.Y., 2020. Metagenomics-based evaluation of groundwater microbial profiles in response to treated wastewater discharge. Environ. Res. 180, 108835. 10.1016/j.envres.2019.108835

Zhang, A.N., Gaston, J.M., Dai, C.L., Zhao, S., Poyet, M., Groussin, M., Yin, X., Li, L.G., van Loosdrecht, M.C.M., Topp, E., Gillings, M.R., Hanage, W.P., Tiedje, J.M., Moniz, K., Alm, E.J., Zhang, T., 2021. An omics-based framework for assessing the health risk of antimicrobial resistance genes. Nat. Commun. 12, 1–11. 10.1038/s41467-021-25096-3

Zhang, J., Buhe, C., Yu, D., Zhong, H., Wei, Y., 2020. Ammonia stress reduces antibiotic efflux but enriches horizontal gene transfer of antibiotic resistance genes in anaerobic digestion. Bioresour. Technol. 295, 122191. 10.1016/j.biortech.2019.122191

Zhang, Yuan, Zhang, Yuanzhu, Kuang, Z., Xu, J., Li, C., Li, Y., Jiang, Y., Xie, J., 2019. Comparison of Microbiomes and Resistomes in Two Karst Groundwater Sites in Chongqing, China. Groundwater 57, 807–818. 10.1111/gwat.12924

